# *In silico* and empirical evaluation of twelve COI & 16S metabarcoding primer sets for insectivorous diet analyses

**DOI:** 10.1101/742874

**Authors:** Orianne Tournayre, Maxime Leuchtmann, Ondine Filippi-Codaccioni, Marine Trillat, Sylvain Piry, Dominique Pontier, Nathalie Charbonnel, Maxime Galan

**Affiliations:** CBGP, INRA, CIRAD, IRD, Montpellier SupAgro, Université de Montpellier, Montpellier, France; Nature Environnement, Surgères, France; LabEx ECOFECT « Ecoevolutionary Dynamics of Infectious Diseases », Université de Lyon, Lyon, France; CNRS, Laboratoire de Biométrie et Biologie Évolutive, UMR5558, Université de Lyon, Université Lyon 1, Villeurbanne, France

**Keywords:** Arthropod, bat, environmental DNA (eDNA), predator feeding, high-throughput sequencing

## Abstract

This last decade, environmental DNA metabarcoding approaches have been developed and improved to minimize biological and technical biases; some challenges, however, remain, as the design of primers. Here we have performed a comprehensive assessment of ten COI and two 16S primer sets. We have combined *in silico, in vivo*-mock community of 33 arthropod taxa from 16 orders and guano analyses to identify primer sets that should maximize arthropod detection and taxonomic identification, whilst identifying bat species and minimizing labour time and cost. We have focused on two insectivorous bat species living in mixed-colonies, the greater horseshoe bat (*Rhinolophus ferrumequinum*) and Geoffroy’s bat (*Myotis emarginatus*). We have found that the level of primer degeneracy is the main factor influencing arthropod detection for *in silico* and mock community analyses, while the amplicon length is critical for the detection of arthropods from degraded DNA samples. Our results confirm the importance of performing predator detection and taxonomic identification, simultaneously with arthropod sequencing, as faeces samples can be contaminated by different insectivorous species. Moreover, amplifying bat DNA does not affect the primers’ capacity to detect arthropods. We therefore recommend the systematic simultaneous identification of predator and prey. Finally, we evidenced that one third of the prey occurrences are unreliable and probably not of primary interest in diet studies, which might decrease the relevance of combining several primer sets instead of using one efficient primer set. In conclusion, this study provides general criteria enabling the selection of primers whilst considering different scientific and methodological constraints.

## 1 INTRODUCTION

Environmental DNA (eDNA) metabarcoding, the genetic analysis of environmental samples such as soil, water or faeces, provides a rapid and cost-effective tool for the study of species that are difficult to detect or monitor (Bohmann et al., 2014). This approach allows the simultaneous identification of the taxa composition of environmental samples, without requiring the initial isolation of organisms (Clare, 2014; Taberlet et al., 2012). eDNA metabarcoding is of particular interest for the diet analysis of rare or elusive species, and this approach has been applied to a large spectrum of organisms (Clare et al., 2009; Shehzad et al., 2012; Rytkönen et al., 2019; Kartzinel and Pringle, 2015; Corse et al., 2017). Compared to the traditional microscopic study of the undigested prey fragments in faecal remains, eDNA metabarcoding has two key advantages for diet analysis: (i) a finer resolution (potentially to the species level), and (ii) the simultaneous processing and sequencing of several hundred samples (Galan et al., 2018). However, eDNA metabarcoding is subject to many biological and technical distortions at each step of its process, including fieldwork, laboratory analysis and bio-informatics. Old-established samples, storage in poor conditions, contaminations, PCR inhibitors, PCR stochasticity and chimera development are common biases influencing the reliability of results (for a review see, Alberdi et al., 2019). Many methodological studies have been made to limit some of these biases and to provide some guidelines for the elaboration of metabarcoding protocols, as for example the systematic inclusion of technical replicates and negative controls (Alberdi et al., 2018; Corse et al., 2017; Elbrecht and Steinke, 2019; Galan et al., 2016; Mata et al., 2018).

Another major issue of eDNA metabarcoding that still needs to be solved, despite a growing attention, is the selection of the primer set(s). This is crucial because primers targeting the DNA fragment to be amplified and sequenced, should be suitable for all the taxa present in the environmental samples. The targeted DNA fragment has to be variable enough to discriminate close species, but also sufficiently abundantly referenced in public sequence databases to allow for the identification of sequences gathered. The cytochrome c oxidase I (COI) mitochondrial gene fulfils these criteria. It is the most widely used gene in animal metabarcoding analyses (Andújar et al., 2018; Hebert et al., 2003). The targeted ‘Folmer region’ of the COI sequence used for the standardized DNA barcoding is 658 base pairs (bp), which is too long to be sequenced efficiently by the second generation of High-Throughput Sequencing (HTS) platforms. Moreover, eDNA is often highly degraded, because of digestion processes (faecal samples), or exposure of samples to the outer environment (*i.e.* rain, sunlight, etc, Oehm et al., 2011). This makes the use of the entire ‘Folmer region’ COI sequence irrelevant (Deagle et al., 2006). ‘Mini COI barcodes’ (*i.e.* with a targeted region range of < 200bp; Hajibabaei et al., 2006; Pompanon et al., 2012) are therefore more commonly used in eDNA metabarcoding analyses, although lack of conserved regions in the COI sequence can make the design of universal primers difficult (Deagle et al., 2014). Thus, some authors have argued for the joint use of several COI primer sets (Corse et al., 2019; Esnaola et al., 2018), or of the combination of mitochondrial 16S rRNA and COI primer sets (Alberdi et al., 2018; Bohmann et al., 2018). This should allow the coverage of the taxonomic spectrum of prey to be increased. For example, Esnaola et al. (2018) have shown that the combination of the Zeale et al. (2011) and Gillet et al. (2015) primer sets allows 37.2% more species to be detected than Gillet’s primer set alone. However, the combination of several primer sets greatly increases not only the financial cost but also the duration of both laboratory work and analyses of metabarcoding studies. In this context, the use of a single highly degenerated primer set is potentially very promising. Some studies have highlighted the efficiency of degenerated primers (Elbrecht and Leese, 2017; Galan et al., 2018; Vamos et al., 2017). Degenerated bases are used in primer’s sequences to avoid mismatches at the variable positions between the different targeted taxa. In theory, this enables the amplification and identification of different taxa with a unique primer set in a single PCR reaction. However, degenerated primers have to be carefully designed because high levels of degeneracy can lead to high rates of non-target amplification (Innis et al., 2012). Up to now, no consensus has been reached and a multitude of primer sets have been designed for the COI and 16S genes that differ in their target length, degeneracy levels and position.

Designing metabarcoding protocols for the study of insectivorous bat diet is still an important issue. Such molecular analysis is critical because the direct observation of bat feeding is virtually impossible. The morphological identification of prey remains in the guano is not resolutive and can be highly time consuming. Finally, many bat species are endangered and therefore protected so that invasive methods cannot be applied to carry out diet surveys. Metabarcoding analysis of the DNA contained in bat guano has thus been developed, (i) to better understand bat ecology (Arrizabalaga-Escudero et al., 2015; Clare et al., 2014a), (ii) to highlight the potential ecosystem services provided by bats as pest suppressors (Aizpurua et al., 2018; Maslo et al., 2017), and (iii) ultimately to set up effective conservation strategies (Arrizabalaga-Escudero et al., 2015). However, most of these studies have used arthropod specific primer sets. The resulting problem when working with guano collected from roost sites comes from the fact that several bat species may roost in the same sites, and that the guano is not easily distinguishable between bat species. In this case, it is critical to identify bat species to avoid mis-assigning preys to the wrong bat species and also to discard guano samples that could be contaminated with excreta from other bat species. To this end, Galan et al. (2018) optimized a metabarcoding approach to simultaneously identify bat species and their prey, instead of using different primer sets and methodologies to identify bat species on one hand and the arthropods on the other hand (e.g. Bohmann et al., 2011; Van den Bussche et al., 2016). As the amplification of bat DNA may reduce the sensitivity of prey DNA detection (Pompanon et al., 2012), it is important to find a trade-off between the success of an exhaustive prey amplification and bat species identification.

In this study, we have addressed this question of the design of an optimal DNA metabarcoding approach in order to study the diet of two European insectivorous bat species: the greater horseshoe bat (*Rhinolophus ferrumequinum*) and the Geoffroy’s bat (*Myotis emarginatus*). These two species are of particular interest regarding this methodological issue because they share maternity roosts during summer but have contrasted diets. It is therefore important to be able to assign guano collected from one or other of these two species. In this context, our main objective was to compare different primer sets to identify the primer characteristics that would maximize the accuracy of arthropod detection and identification, while minimizing the time and cost of labour. We also compared the capacity of primer sets to provide identification of bat species without over amplifying bat DNA. Following the recent recommendations of Elbrecht et al. (2019), we carried out a complete primer assessment consisting of three stages. We first made an *in silico* comparison of primer sets, based on hundreds of thousands of arthropods’ sequences gathered from public databases. This step enabled to evaluate the primers’ efficiency to detect a wide taxonomic array of arthropods, independently of the quality of the samples and the effects of laboratory procedures (extraction, PCR, etc). We then performed an *in vivo* comparison of primer sets on two mock communities (MC) – one containing only arthropod DNA and the other one containing both arthropod and bat DNA. This step enabled to evaluate the efficiency of primer sets in detecting a wide taxonomic array of known arthropod taxa, and to assess whether the presence of bat DNA influences the efficiency of detecting prey. Lastly, we made an *in vivo* comparison of primer sets on guano samples from two insectivorous bat species. This step allowed to compare the efficiency of primer sets in amplifying degraded arthropod DNA which is likely to be encountered when working with guano samples collected from roost sites.

## 2 MATERIALS AND METHODS

### 2.1 *In silico* evaluation

First, we selected primers commonly used for arthropod detection and identification in metabarcoding approaches (Table 1). Six primer sets were selected on the COI gene and one on the 16S gene. We favoured primer sets targeting either short region length (between 133 and 218 bp), but also some longer fragments (between 313 and 322 bp) to evaluate the potential effect of primer length on prey detection in bat guano. Then, based on existing primers, we designed four new COI combinations of primers and one new 16S combination of primers on the basis of the seven previously chosen primer sets. For the COI reverse primer from Jusino et al. (2018) and the 16S primers from Epp et al. (2012), we increased the base degeneracy level to improve their success of hybridization on DNA prey during the PCR amplification. To estimate the efficiency of the 12 primer sets, we applied the R package *PrimerMiner* (Elbrecht and Leese, 2016) to cluster 4,259,845 sequences from BOLD database in 327,412 COI OTUs (Operational Taxonomic Units) and 83,651 sequences from NCBI database in 25,505 16S OTUs for 21 arthropod orders including potential bat prey (details in Table S1). COI and 16S sequences were aligned separately using MAFFT v7.017 (Katoh et al., 2002) as implemented in GENEIOUS v8.1.7 (Kearse et al., 2012). The consensus sequence alignment for each order of arthropods was visualized using *PrimerMiner* which allows to identify suitable primer binding sites (Figure S1). All primer sets are detailed in Table 1. They varied in target region length and degeneration level (Figure 1).

**Table 1.**
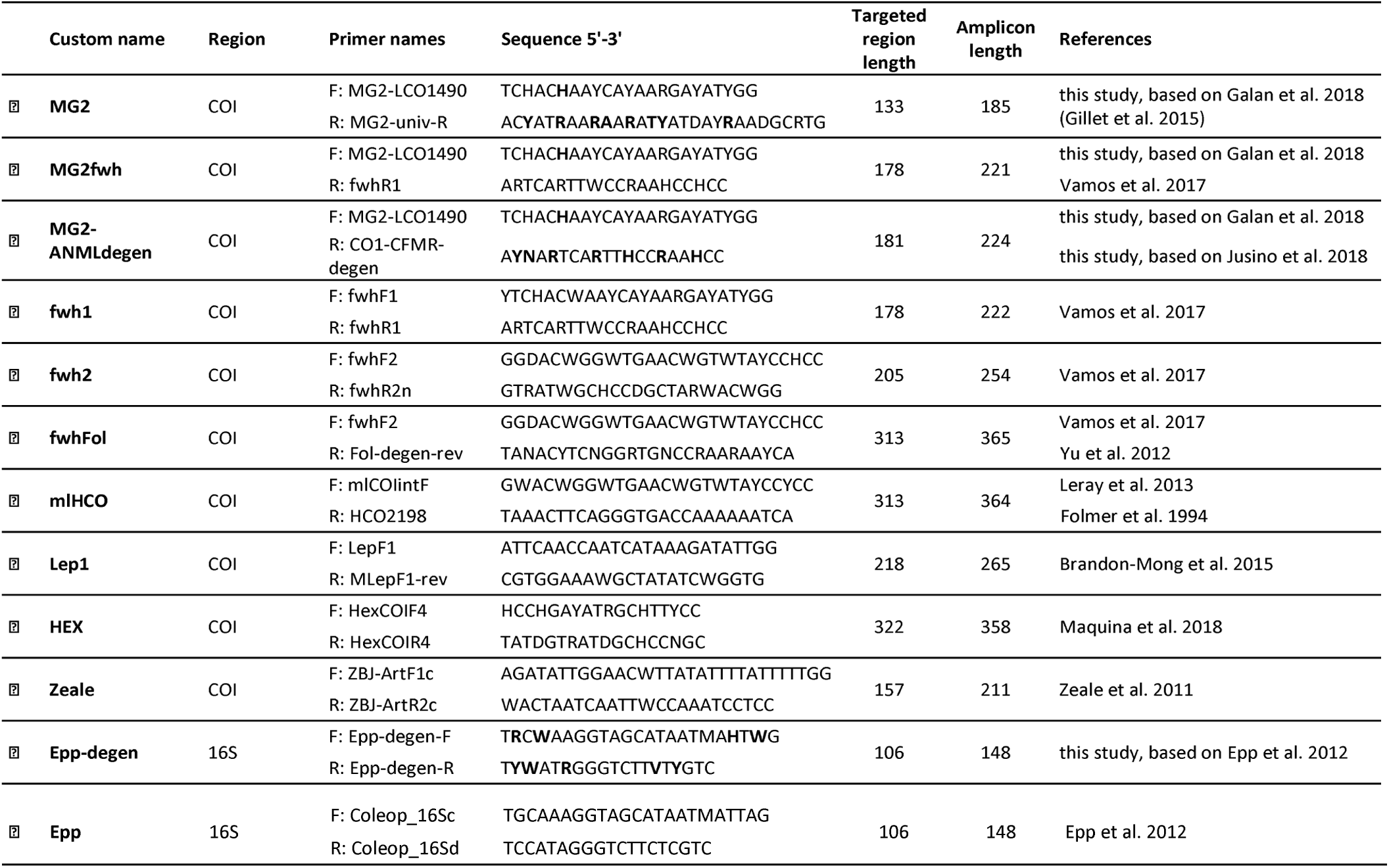
Primer sets information. Bases in bold indicate the bases which were degenerated in this study.

**Figure 1.**
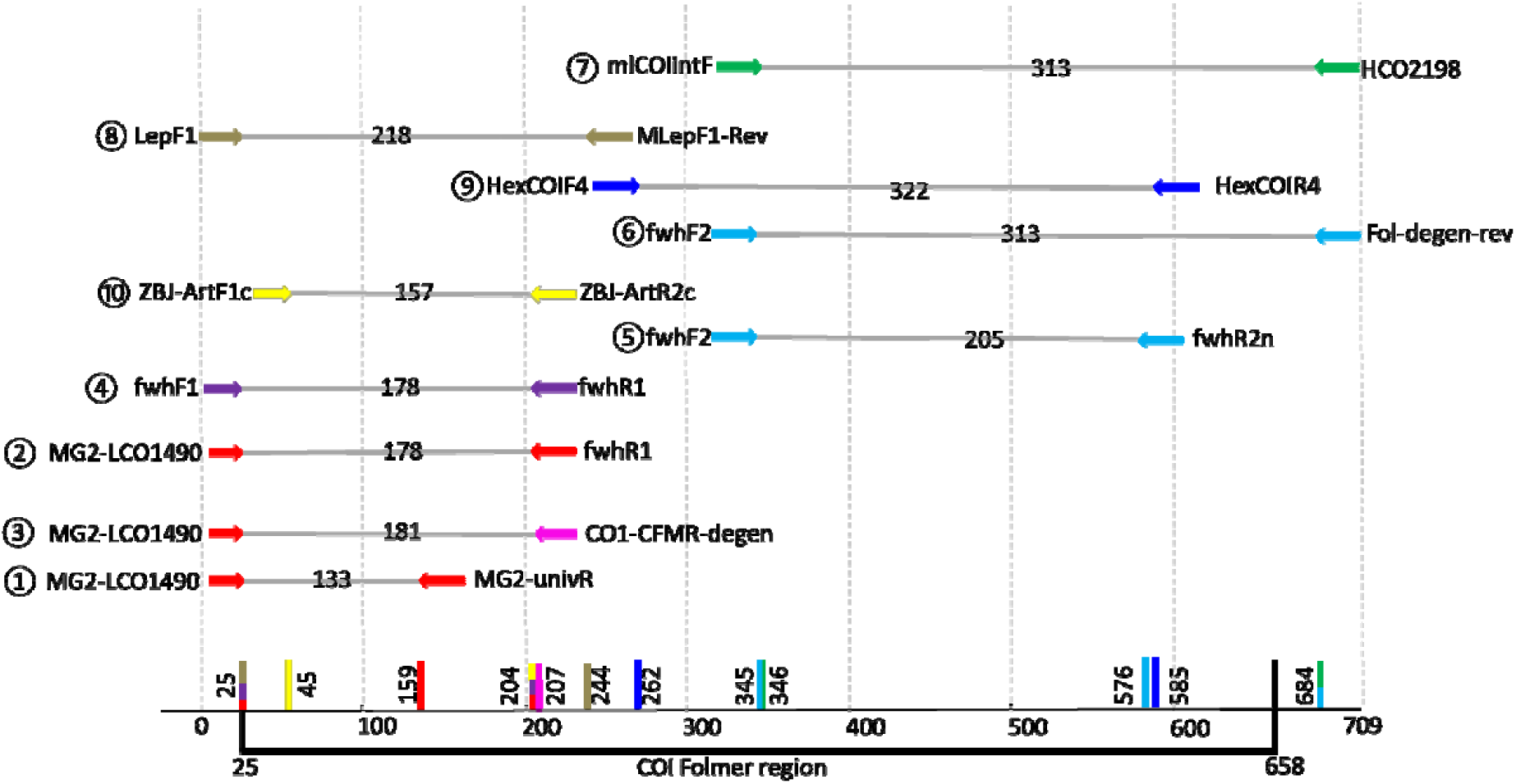
Visual representation of primer length and position on the COI gene. Primers are represented by a number (see Table 1) and a coloured arrow, with each colour representing a unique primer set. For each primer set, the number on the grey line correspond to the amplicon length. This information is synthetized on the bottom of the figure, with the whole COI Folmer region represented and each primer set indicated with traits of its respective colour.

These 12 primer sets were evaluated and compared using *PrimerMiner*. This comparison is based on a mean penalty score per arthropod order. It is calculated as a mismatch scoring that considers the adjacency, position and type of mismatch between primers and template sequences. Primers’ evaluation was only made for arthropod orders represented by at least 100 OTUs, as recommended by Elbrecht and Leese (2016). This enables to capture a sufficient part of the variability potentially existing at primer binding sites. We evaluated each primer set by combining the penalty scores from both the forward and reverse primers. We determined whether the arthropod OTUs should be theoretically successfully amplified using the default value (success: penalty score < 120).

### 2.2 *In vivo* evaluation

#### 2.2.1 Mock community: sample collection and preparation

Sampling details are provided in Table S2. Briefly, we collected 33 arthropod individuals representing 16 orders (two taxa per order except for Opiliones, Dermaptera, Isopoda, Psocodea, Julida, Polyxenida and Raphidioptera). They were captured alive in May and June 2019 in the South of France, then immediately frozen at −20°C to avoid DNA degradation. Bat DNA was obtained from tissue samples. They were collected in Western France from the wing membrane (patagium) of greater horseshoe bats (*R. ferrumequinum*) using a 3-mm diameter biopsy punch. Bat samples were preserved in 95% ethanol solution at 4°C until DNA extraction and pooling.

The DNA of arthropods and bats was extracted using EZ-10 Spin Column Genomic DNA Minipreps Kit for Animal (BioBasic) following the manufacturer’s instruction. DNA extractions were normalized to 7 ng/µL after Qubit fluorimeter quantification (Invitrogen). The integrity of DNA was evaluated by electrophoresis on a 1,5% agarose gel.

We built two versions of the same mock community (MC) of 33 different arthropods taxa. The first one contained the 33 arthropod DNA extracts mixed in equimolar proportion (MC_arthr_). The second one contained 50% of this mock community and 50% of bat DNA (MC_arthr+bat_).

We used the same DNA extractions of bats and arthropods to build reference sequences for the COI and 16S genes. Normalized DNA was amplified and sequenced for each individual to provide reference sequences for each gene. Firstly, we sequenced the 658bp ‘Folmer region’ of the COI gene using Sanger technology as described in Sow et al. (2018). Secondly, we sequenced the 106bp target region of the two 16S minibarcodes using the Epp-degen primer set and MiSeq sequencing technology for each DNA extractions independently, following the same protocol used for the analysis of the mock and guano samples, as described below.

#### 2.2.2 Guano samples: collection and preparation

We collected 22 faecal pellets from five mixed maternity colonies of the greater horseshoe bat and Geoffroy’s bat in Western France (see details in Table S2). Each guano was retrieved from paper plates let on the ground during 10 days using single-usage pliers in order to avoid contaminations between samples. Paper plates were renewed at each collection to avoid contaminations. Samples were stored at −20°C until DNA extraction.

DNA extraction was performed according to Zarzoso-Lacoste et al. (2018). Briefly, guano samples were frozen at −80°C then bead-beaten for 2 x 30s at 30 Hz on a TissueLyser (Qiagen) using a 5-mm stainless steel bead. DNA was extracted using the NucleoSpin 8 Plant II kit (Macherey Nagel) with the slight modifications recommended in Zarzoso-Lacoste et al. (2018).

#### 2.2.3 PCR and library construction

We made four versions of each primer by adding a 5’ heterogeneity spacer of 0 to 3 bp to the primer sequence (Table S3). This increased the diversity at each sequencing cycle, improved the detection of the sequencing clusters at the flowcell surface and thus increased the quality of the reads. The four versions of each primer were mixed together before PCR.

We used the two-step PCR protocol described in Galan et al. (2018) with slight modifications. First, we increased the time of extension step (2min instead 45s) in PCR_1_ and PCR_2_ to reduce chimera formation. Second, for each PCR replicate multiplexed in the same run, we used a double indexing strategy as recommended in Kircher et al. (2012): each 9-bp i5 and i7 dual-index were used only for one PCR sample, eliminating the problem of ‘leak’ due to false index-pairing (Martin, 2019). The lists of the dual-index used are described in Table S4. We used the same PCR programs for all the primer sets with a low annealing temperature (45°C) as recommended in recent studies (Elbrecht et al., 2019; Jusino et al., 2018). We validated this program by checking the quality of PCR amplification on the mock community for the 12 primer sets (Figure S2). The PCR_1_ was performed in 10 µl reaction volume using 5 µl of 2x Qiagen Multiplex Kit Master Mix (Qiagen), 2.5 µl of ultrapure water, 0.5 µl of each mix of forward and reverse primers (10 µM), and 1.5 µl of DNA extract. The PCR_1_ conditions consisted in an initial denaturation step at 95°C for 15 min, followed by 40 cycles of denaturation at 94°C for 30 s, annealing at 45°C for 45 s, and extension at 72°C for 2min, followed by a final extension step at 72°C for 10 min. The PCR_2_ consisted in a limited-cycle amplification step to add multiplexing index i5 and i7 (9 bases each) and Illumina sequencing adapters P5 and P7 at both ends of each DNA fragment from PCR_1_. PCR_2_ was carried out in a 10µl reaction volume using 5 µl of Qiagen Multiplex Kit Master Mix (Qiagen) and 2µL of each indexed primer i5 and i7 (0.7 µM). Then, 2 µL of PCR_1_ product was added to each well. The PCR_2_ started by an initial denaturation step of 95°C for 15 min, followed by 8 cycles of denaturation at 94°C for 40 s, annealing at 55°C for 45 s and extension at 72°C for 2min followed by a final extension step at 72°C for 10 min.

We included in each 96-well microplate a negative control for extraction (NC_ext_), a negative control for PCR (NC_PCR_) and a negative control for indexing (NC_index_). We performed three PCR technical replicates per sample on each DNA extract. For the guano samples, we considered a positive sample for a particular taxon if at least two replicates over three were positive, to overcome the stochasticity of the PCR in the detection of rare prey while reducing the number of putative false-positive results (*i.e.* one positive replicate over three is considered as a false-positive result, Alberdi et al., 2018).

We checked the homogeneity of amplifications between primer sets and the absence of nonspecific amplification by electrophoresis of 3 µL of each PCR_2_ product on a 1.5% agarose gel. Then PCR_2_ products were pooled separately for each of the 12 primer sets and put on a low-melting agarose gel (1.25%) for excision. After electrophoresis, the excision step aimed at eliminating the primer dimers and non-specific amplifications. We used the PCR Clean-up Gel Extraction kit (Macherey-Nagel) to purify the excised bands. The 12 pools were quantified using the KAPA library quantification kit (KAPA Biosystems) taking into account the different fragment length, normalized at 4nM, and pooled in equimolar concentration before loading 14 pM and 5% of PhiX control on a MiSeq flow cell with a 500-cycle Reagent Kit v2 (Illumina).

#### 2.2.4 Taxonomic assignments

First, we used a R pre-processing script (Sow et al., 2019) to merge pair sequences into contigs with FLASH v.1.2.11 (Magoč and Salzberg, 2011) and to trim primers with CUTADAPT v.1.9.1 (Martin, 2011). We then used the FROGS pipeline (‘Find Rapidly OTU with Galaxy Solution’, Escudié et al., 2018) to create an abundance table for each variant. Briefly, this pipeline enabled to (i) filter sequences by length (+/-20 bp from the expected length), (ii) cluster in Operational Taxonomic Units (OTUs) the variants using a maximum aggregation distance of one mutation with the SWARM algorithm (Mahé et al., 2014), (iii) remove chimeric variants using VSEARCH with *de novo* UCHIME method (Edgar et al., 2011; Rognes et al., 2016), and (iv) filter by keeping only OTUs present in at least two PCR replicates.

Taxonomic assignments were carried out for each primer set, following different procedures. We used the reference sequences produced to analyse mock community results (see above). This enabled to identify the genuine sequences of the two mock communities (Table S5). With regard to guano samples, we analysed the 16S OTUs using BLASTN (Altschul et al., 1990) and the NCBI Nucleotide database (Benson et al., 2008). Taxonomic assignments of COI OTUs were made using the NCBI BLAST+ automatic affiliation tool available in the FROGS pipeline, with the Public Record Barcode Database (data related to BOLD database http://v3.boldsystems.org in February 2019, with maximum 1% of N). We determined the final affiliations following the description provided in Appendix S1.

### 2.3 Statistical analyses

#### 2.3.1 In silico data

We tested the effect of the level of primer degeneracy on the mean penalty score and on theoretical amplification success. We used respectively a Poisson and a binomial Generalized Linear Mixed Model (GLMM), with the primer set and the order as random effects.

#### 2.3.2 Mock communities data

We tested the effect of the amplicon length, the level of primer degeneracy and the percentage of bat reads on the percentage of arthropod taxa detected using a binomial Generalized Linear Model (GLM).

#### 2.3.3 Guano samples data

We tested the effect of the amplicon length, the level of primers degeneracy, the total number of reads and the percentage of bat reads on the number of arthropod occurrences using a Poisson GLM. Taxa occurrence was considered instead of the absolute number of taxa in order to take into account the frequency of detection of a particular taxon for each primer set, and to minimize the detection of rare taxa (*i.e.* detected in a single sample).

We did not include the 16S primer sets in the *in vivo* statistical analyses (mock communities and guano samples) because of confounding factors (e.g. smaller size of the 16S reference database compared to the COI database).

#### 2.3.4 Defining a strategy to determine the best primer set(s) for the study

We developed a multi-criteria table that included criteria for each step of the primer set evaluation (*in silico*, mock community and guano sample analyses). We provided a score for each criteria with regard to the objectives and constraints of our future metabarcoding studies. All criteria are detailed in Table S7.

## 3 RESULTS

### 3.1 *In silico* evaluation

The *in silico* evaluation was performed on 21 arthropod orders (see list in Table S1). We recovered almost fifty times more sequences for the COI (4,259,845 sequences) than for the 16S gene (83651 sequences) using BOLD and NCBI databases. The two orders Dermaptera and Julida, and the four orders Archaeognatha, Dermaptera, Julida and Mecoptera, were respectively excluded from the COI and 16S analyses using *PrimerMiner* because of their insufficient number of OTUs (< 100; Table S1).

The mean penalty scores and the theoretical amplification success varied strongly between primer sets and between arthropod orders (Figure 2, Tables S8 and S9). We found a significant negative influence of the level of primer degeneracy on the mean penalty scores (GLMM, *p* = 1.01e-12) and a positive influence on the theoretical amplification success (GLMM, *p* = 2.88e-09) (Table S10). The high mean penalty scores (> 165) and low amplification success (mean < 50%) were observed for the primer sets exhibiting a lack of degeneracy (e.g. mlHCO, Lep1, Zeale and Epp). For example, the mean penalty score of Epp primers was around 12 times higher than the score of its degenerated version Epp-degen.

**Figure 2.**
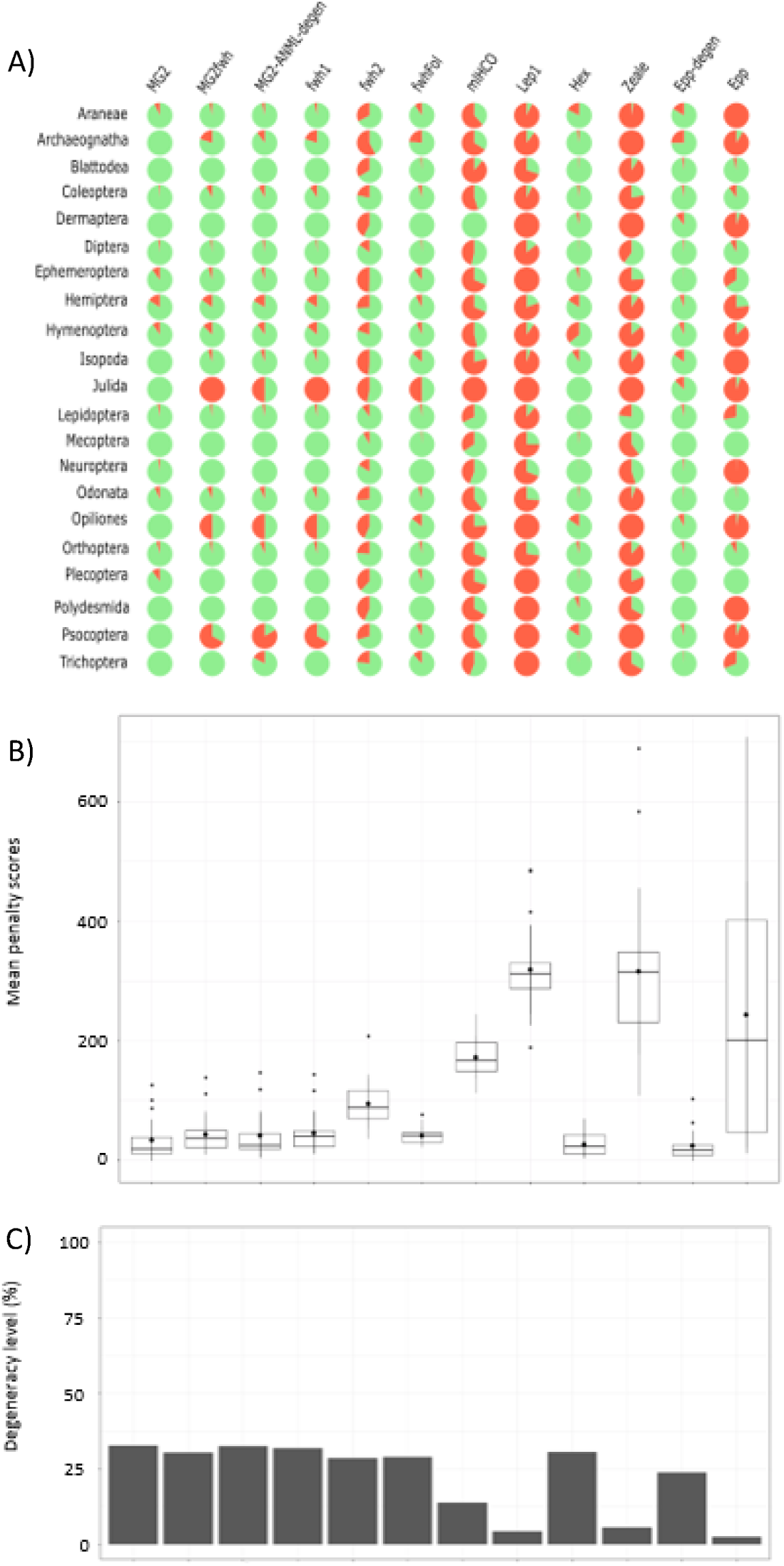
*In silico* evaluation of arthropod orders represented by at least 100 OTUs using *PrimerMiner.* A) Primer sets performance is shown for each order using pie charts, with green and red colours representing respectively success and failure of amplification. Success of amplification corresponded to *PrimerMiner* mean penalty score < 120 and amplification failure to a mean penalty score >= 120. B) Boxplots of the median of *PrimerMiner* mean penalty scores over all arthropod orders and for each primer set, with mean values represented by a circle within boxplots. C) Percentage of degeneracy level of each primer set.

### 3.2 *In vivo* evaluation - Mock communities

#### 3.2.1 Sequencing results

DNA quality checks revealed that DNA showed a good integrity for all the taxa considered, except for the Hemiptera-1 (*Uroleucon sp.*) which was degraded (Figure S2). PCR products of the MCs on agarose gels revealed that amplification was homogeneous between primer sets (Figure S2). MiSeq sequencing produced a total of 3,100,343 reads for the two MCs. Removing reads with unexpected length excluded up to 6.3 % of the reads (Hex primer set), removing chimera excluded up to 4.67% of the reads (fwhFol primer set) and removing reads not shared by at least two PCR replicates excluded up to 21.64% of the reads (mlHCO primer set). The remaining reads varied from 168,815 for Hex to 284,216 for Lep1 (Table S11).

#### 3.2.2 Detection of bat DNA

All primer sets amplified less bat DNA (up to 18.36% of the number of reads; replicate 2 of Epp-degen) than expected (50% because arthropod and bat DNA were in equimolar proportions in the MC_arthr+bat_, see Figure S3). Primers with a lack of degeneracy amplified only few reads (Hex, mlHCO and Epp; mean of triplicate < 0.01%) or did not amplify any bat DNA at all (Zeale and Lep1). The degenerated primers fwh2 and MG2-ANML-degen amplified less than 1% of bat reads and fwh1 and MG2fwh less than 5.2%. The best primer sets for the amplification of bat DNA were Epp-degen (mean triplicate 10,17%), fwhFol (mean triplicate 7.53%) and MG2 (mean triplicate 7.43%).

#### 3.2.3 Detection of arthropod taxa

The percentage of arthropod taxa detected in MCs varied from 67% to 100% (see Figure 3 and Figure S4). In the mock community with no bat DNA (MC_arthr_), only MG2 amplified all arthropod orders in triplicate. Five other primer sets (MG2fwh, MG2-ANML-degen, fwh1, fwh2 and fwhFol) amplified all arthropod orders; however, the taxa with degraded DNA (*Uroleucon sp.* Hemiptera-1) amplified only few reads in two replicates over three. Hex was also among the best primer sets as it amplified all taxa except the degraded Hemiptera-1. Hemiptera-1 was mis-amplified by less than half of the primer sets (only one to three reads were recorded) and not amplified by the others. In accordance with the *in silico* evaluation, mlHCO, Lep1, Zeale and Epp failed to detect an important number of taxa, and Zeale and Lep1 were the less efficient primer sets with respectively eight and 14 taxa not amplified or mis-amplified (two replicates over three). Our results revealed a positive effect of the COI primers degeneracy level on the percentage of detected taxa in MC_arthr_ (GLM, *p* = 6.54e-05; Table S10) but no effect of the amplicon length (GLM, *p* = 0.681; Table S10). The influence of the degeneracy is also illustrated by the two 16S primer sets: Epp mis-amplified or did not amplified five (MC_arthr_) and six taxa (MC_arthr+bat_), while its degenerated version Epp-degen mis-amplified or did not amplify only two taxa over the 33 of each mock community.

**Figure 3.**
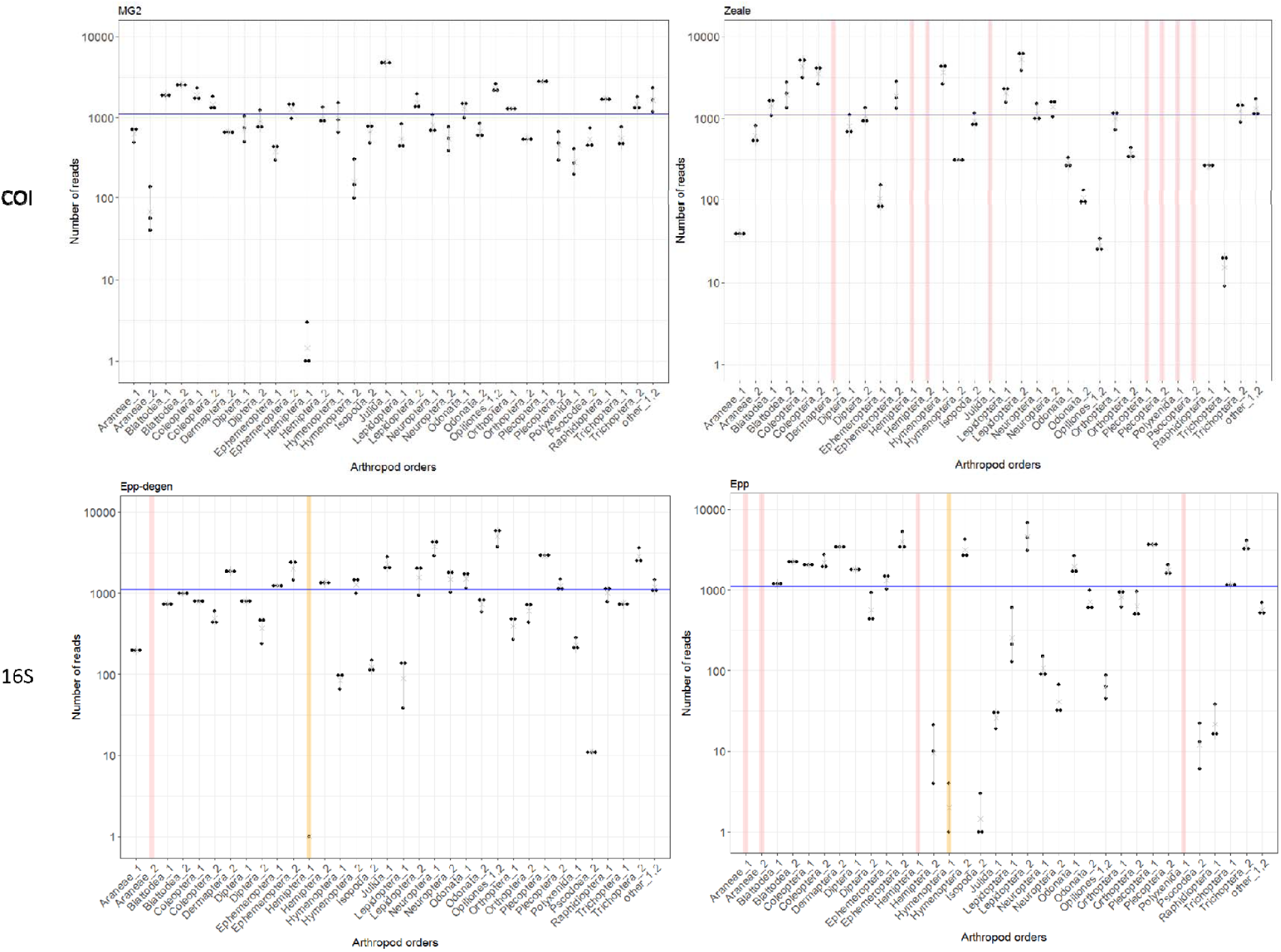
Representation on a log-scale of the number of reads gathered for each arthropod order with the MG2 and Zeale COI primer sets, and Epp and Epp-degen 16S primer sets, from the three technical replicates of the mock community MC_arthr_. The blue line indicates the expected number of reads. Each point represents a technical replicate. Yellow bars emphasize situations where only one or two PCR over three amplified. Red bars emphasize situations where none of the replicate amplified.

In the mock community with bat DNA (MC_arthr+bat_), four primer sets MG2, MG2-ANML-degen, fwh2 and fwhFol amplified all taxa in triplicates (Figure S4). The primer set fwh1 performed with reduced efficiency as it did not amplify Hemiptera-1 at all. The two primer sets Zeale and Lep1 were still the less efficient ones with respectively eight and 13 taxa that were not amplified or mis-amplified. Our results revealed a positive effect of the level of COI primers degeneracy on the percentage of detected taxa in MC_arthr+bat_ (GLM, *p* = 0.001; Table S10). We found no effect of the amplicon length (GLM, *p* = 0.781; Table S10) or of the percentage of bat reads (GLM, *p* = 0.668; Table S10).

In both MCs, the primer sets that detected the smaller number of taxa also exhibited the largest variation of read number between arthropod taxa and between the observed and expected number of reads (Zeale, Lep1, mlHCO and Epp, Figure S4).

#### 3.2.4 Taxonomic identification of arthropod taxa

The COI primer sets provided identical identification and exhibited only slight differences in taxonomic resolution (Table S12). By contrast, the COI and 16S primer sets often provided different identifications and levels of taxonomic resolution. Identical identifications were found for only eight taxa among the 33 arthropod taxa included in the MCs. The 16S primer sets always reached lower level of taxonomic resolution. The only exceptions were (i) Lepidoptera-2 that was identified at the species level (*Melitae deione*) by the 16S primer sets and at the genus level only by the COI primer sets, and (ii) Psocoptera-2 that was identified at the genus level (*Myopsocus sp.*) by the 16S primer sets and not identified by the COI primer sets.

### 3.3 *In vivo* evaluation – Guano samples

#### 3.3.1 PCR verification and sequencing results

DNA amplification was relatively homogeneous between primer sets and pellet samples, except for Hex and Lep1 which led to amplification failures and non-specific products (Figure S2). MiSeq sequencing produced a total of 9,190,350 reads. Removing reads with unexpected length excluded 0.52% (Epp-degen) to 10.57% (Lep1) of the reads (except 40.16 % of the reads for Hex). Removing chimera excluded 0.1% (Epp-degen) to 6.42% (mlHCO) of the reads. Removing reads not shared by at least two PCR replicates excluded 0.3% (Epp-degen) to 16.17% (Hex) of the reads (Table S11). Finally, the primer set Hex apart (368,121 reads), the remaining reads varied from 552,914 (Epp-degen primer set) to 808,594 (Zeale primer set).

#### 3.3.2 Detection of bat taxa

Two bat species were mainly identified: *Rhinolophus ferrumequinum* and *Myotis emarginatus*. *R. ferrumequinum* was predominant (mean percentage of reads > 98.9% of all the Chiroptera reads) in ten samples, *M. emarginatus* in eleven samples (mean percentage of reads > 99.2%) and both species were found in high mixed proportions in one sample (mean percentage of *R. ferrumequinum* reads = 84.2% and mean percentage of *M. emarginatus* reads = 15.80%). This sample was discarded from further analyses. For eight primer sets and 12 samples, a mix of *R. ferrumequinum* and *M. emarginatus* reads was also observed but in highly unbalanced proportion, indicating very slight traces of cross-contaminations between pellets from different bat species in the colonies (median of reads contamination < 0.05%, Table S13). Traces of DNA from another bat species, *Myotis myotis*, were also found in three samples by four primer sets (mean of 0.0001% of bat reads; Epp-degen, fwh1, MG2 and MG2fwh). Moreover, some primer sets amplified none of the two bat species (Hex) or only one bat species (Zeale, Lep1) whereas others amplified both bat species (Figure S5). However, the percentage of bat reads in *R. ferrumequinum* pellets was always lower than in *M. emarginatus* pellets, whatever the primer set considered (Figure S5).

#### 3.3.3 Detection of arthropod taxa

We found a negative effect of the amplicon length (*p* = 0.030) and a positive effect of the number of reads (*p* = 0.032) on the number of occurrences detected (Table S10). However, the number of reads was not significant (*p* = 0.397) when the primer set with the lowest number of reads (Hex) was excluded. The degeneracy level of the primers and the percentage of bat reads had no effect on the number of occurrences detected (*p* = 0.140 and *p* = 0.111, respectively).

Taken together, our result showed that the 12 primer sets allowed the identification of 96 occurrences of prey items in the *R. ferrumequinum* samples (10 orders, 32 families, 56 genus and 60 species) and 109 occurrences of prey items in the *M. emarginatus* samples (11 orders, 34 families, 63 genus and 65 species). The 16S primer sets revealed respectively about 30% and 20% of the occurrences in *R. ferrumequinum* and *M. emarginatus* samples (Figure 4A). Only six of the ten COI primer sets allowed the detection of at least 50% of the occurrences of prey items in *R. ferrumequinum* samples and only four in *M. emarginatus* samples (Figure 4A). The other COI primer sets revealed from 31.2% to 49.5% of the total number of occurrences. Zeale primer set revealed the highest number of arthropod occurrences for both bat species (N = 68; 62% of *R. ferrumequinum* occurrences and N= 68, 71% of *M. emarginatus* occurrences); however, it did not allow the amplification and identification of bat species. The optimal primer set that amplified bat DNA and provided the highest number of arthropod occurrences for both bat species was fwh1 (number of occurrence_(R.ferrumequinum)_ = 60; number of occurrence_(M.emarginatus)_ = 62) although MG2 was slightly better than fwh1 for *R. ferrumequinum* samples (number of occurrence_(R.ferrumequinum)_ = 63). The combination of one to four primer sets (Epp-degen, fwh1, MG2 and Zeale) allowed a gain from nine to 18 occurrences in *R. ferrumequinum* samples (89.58% of the occurrences detected by combining all primer sets) and from 20 to 25 occurrences in *M. emarginatus* samples (85.32% of the occurrences detected by combining all primer sets) (Figure 4B). However, one third of all the occurrences corresponded to mis-amplification associated with a low number of reads (amplification in two PCRs out of three, Figure 4B and Figure S6). In *R. ferrumequinum* samples, these unreliable occurrences represented a very low number of reads whatever the primer set considered (median < 39 reads). In *M. emarginatus* samples, these unreliable occurrences also represented a very low number of reads ranging from 5 to 575 (except 4,186 for Zeale) for all primer sets (Figure 4B). The maximum number of unreliable reads was lower in *R. ferrumequinum* than in *M. emarginatus* samples (591 reads, Zeale, Figure 4B). Finally, we observed that 79.3% (*M. emarginatus*) and 86.7% (*R. ferrumequinum*) of the occurrences revealed by only one primer set were unreliable (Figure S6).

**Figure 4.**
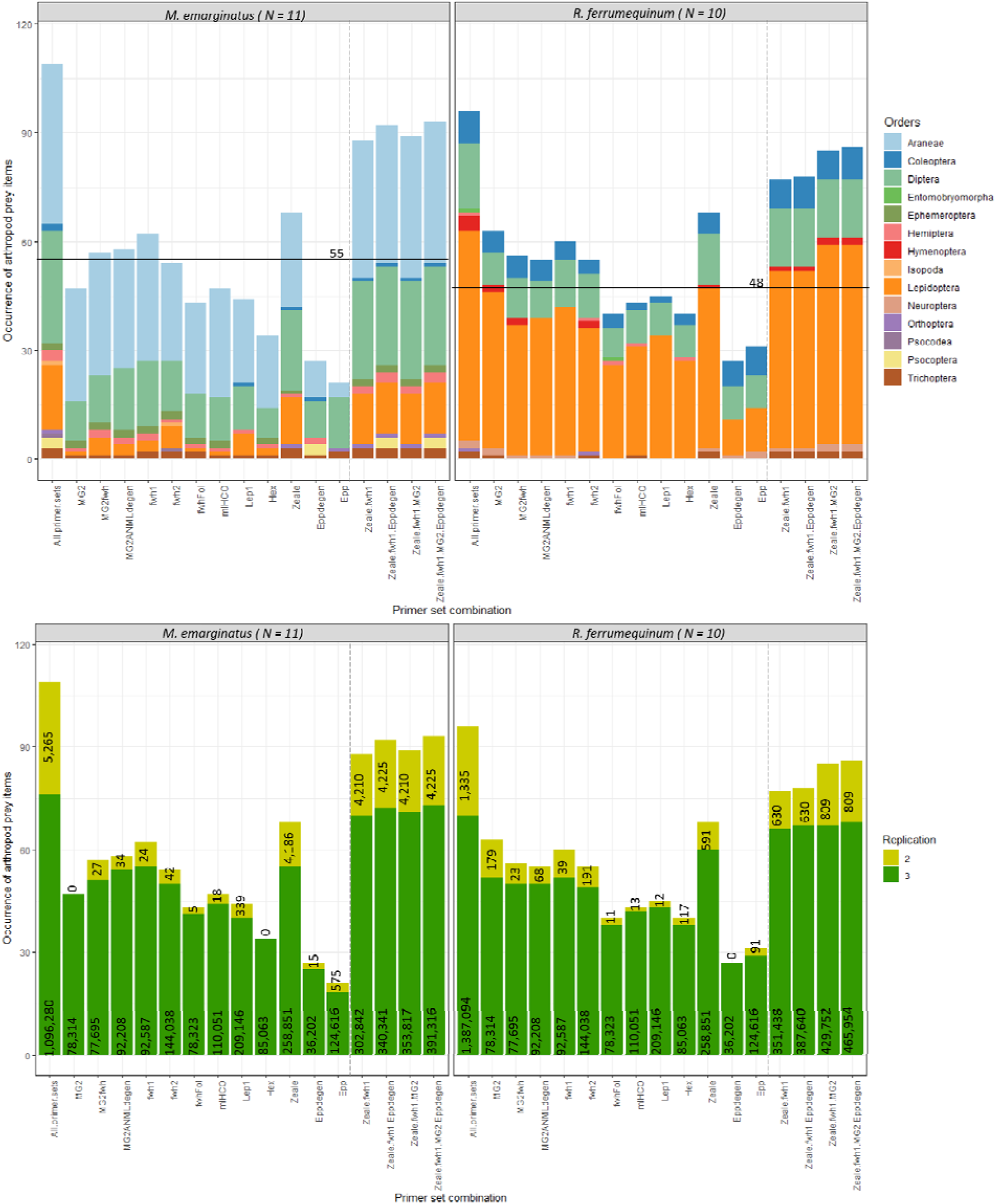
Occurrences of arthropod prey items detected in guano samples for each primer set and for four combination of primer sets. These later include primer sets that provided the best results in terms of occurrence of arthropod orders. Dashed lines separate the primer sets from the combinations of primer sets. N is the number of guano samples analysed for each bat species. A) Occurrences of arthropod orders. Black lines represent 50% of the occurrences (55 occurrences for *M. emarginatus* samples and 48 for *R. ferrumequinum* samples). B) Comparison of the number of occurrences of arthropod orders considering the repeatability of the technical PCR replicates (dark green = occurrences validated in three PCR over three; light green = occurrences validated in two PCR over three). Numbers correspond to the number of reads gathered for each class of occurrence validation (three PCR over three vs two PCR over three).

### 3.4 Multi-criteria evaluation of primer sets

Based on the *PrimerMiner* scores of mean penalty and theoretical amplification success, the multi-criteria table enabled to identify seven appropriate primer sets (MG2, MG2fwh, MG2-ANML-degen, fwh1, fwhFol, Hex and Epp-degen), one intermediate (fwh2) and four inefficient primer sets (Lep1, mlHCO, Zeale and Epp) (Table 2). The number of detected taxa (bat and arthropods) and the taxonomic resolution of their identification indicated that six primer sets were equivalent considering mock community results: MG2, MG2fwh, MG2-ANML-degen, fwh1, fwh2 and fwhFol (Table 2). Finally, Zeale was identified as the optimal primer set when bat DNA amplification is not needed, otherwise fwh1 or MG2 primer sets should be selected (Table 2).

**Table 2.**
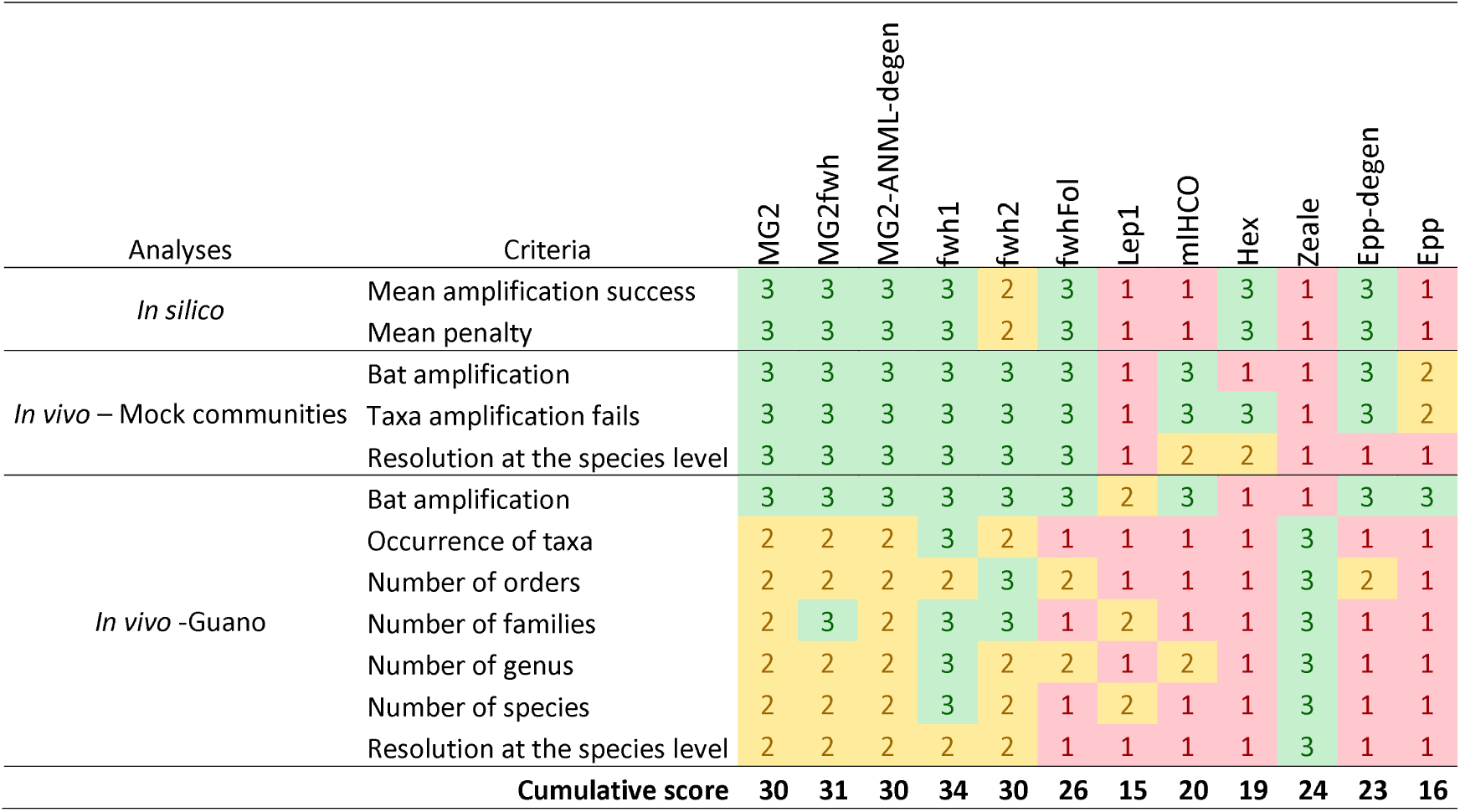
Multi-criteria table indicating the score of each primer set for the three assessment steps performed.

| Analyses | Criteria | MG2 | MG2fwh | MG2-ANML-degen | fwh1 | fwh2 | fwhFol | Lep1 | mHCO | Hex | Zeale | Epp-degen | Epp |
| --- | --- | --- | --- | --- | --- | --- | --- | --- | --- | --- | --- | --- | --- |
| *In silico* | Mean amplification success | 3 | 3 | 3 | 3 | 2 | 3 | 1 | 1 | 3 | 1 | 3 | 1 |
|  | Mean penalty | 3 | 3 | 3 | 3 | 2 | 3 | 1 | 1 | 3 | 1 | 3 | 1 |
| *In vivo* – Mock communities | Bat amplification | 3 | 3 | 3 | 3 | 3 | 3 | 1 | 3 | 1 | 1 | 3 | 2 |
|  | Taxa amplification fails | 3 | 3 | 3 | 3 | 3 | 3 | 1 | 3 | 3 | 1 | 3 | 2 |
|  | Resolution at the species level | 3 | 3 | 3 | 3 | 3 | 3 | 1 | 2 | 2 | 1 | 1 | 1 |
| *In vivo* -Guano | Bat amplification | 3 | 3 | 3 | 3 | 3 | 3 | 2 | 3 | 1 | 1 | 3 | 3 |
|  | Occurrence of taxa | 2 | 2 | 2 | 3 | 2 | 1 | 1 | 1 | 1 | 3 | 1 | 1 |
|  | Number of orders | 2 | 2 | 2 | 2 | 3 | 2 | 1 | 1 | 1 | 3 | 2 | 1 |
|  | Number of families | 2 | 3 | 2 | 3 | 3 | 1 | 2 | 1 | 1 | 3 | 1 | 1 |
|  | Number of genus | 2 | 2 | 2 | 3 | 2 | 2 | 1 | 2 | 1 | 3 | 1 | 1 |
|  | Number of species | 2 | 2 | 2 | 3 | 2 | 1 | 2 | 1 | 1 | 3 | 1 | 1 |
|  | Resolution at the species level | 2 | 2 | 2 | 2 | 2 | 1 | 1 | 1 | 1 | 3 | 1 | 1 |
|  | Cumulative score | 30 | 31 | 30 | 34 | 30 | 26 | 15 | 20 | 19 | 24 | 23 | 16 |

## 4 DISCUSSION

The recent technique of eDNA metabarcoding has proven to be a promising tool for describing biodiversity in a broad array of contexts (Bohmann et al., 2014). Beyond the objectives of scientific quality, the accuracy and completeness of the results gathered by metabarcoding are critical as they are potentially at the core of further management strategies. Previous metabarcoding studies have highlighted how the choice of primer sets may influence the detection of particular arthropod species in diet analyses, and in turn the interpretation of trophic ecology (e.g. for bats, Esnaola et al., 2018).

### 4.1 Need for primers that identify predator species and discard contaminated samples

The requirement for the molecular identification of predators in insectivorous diet analyses depends on the sampling scheme and on the ecology of these organisms. It is especially important when using environmental samples, such as faecal samples, to ensure that samples correspond to the species of interest and to avoid erroneous assignation of prey (Ware et al., 2019). For example, a metabarcoding study by Forin-Wiart et al. (2018) on cat faecal samples revealed that 2.4% of these later belonged to another predator species. Similarly, Biffi et al. (2017) identified the presence of several host species in 57% of the faecal samples supposed to come from the Pyrenean desman. In the particular case of insectivorous bats, the identification of bat species from guano is critical when guano is collected from under the roost, especially when bat colonies are known to be mixed, whereas it is potentially less important when guano is retrieved from trapped bats, or from mono-specific bat colony. However, even colonies supposed to be mono-specific can be shared by cryptic bat species (Filippi-Codaccioni et al., 2018) or other insectivorous species (e.g. birds). Furthermore, we showed that some of the guano sampled under the roosts of mixed colonies (*R. ferrumequinum* / *M. emarginatus*) are contaminated with excreta from different bat species, including *M. myotis* that are not expected to be present in the studied colony. Hence, in future diet analyses, the simultaneous identification of bat species will enable the presence of unexpected species in the roost to be revealed and also allow guano that is too contaminated to be discarded. It is thus particularly important to ensure that primer sets are able to identify all bat species potentially present in the roosts. Indeed, our results showed that some primer sets do not amplify bat species at all, and that others can only provide bat identification for some of the bat species of interest (one out of three).

The simultaneous identification of predator and prey is often avoided in diet analyses as an elevated predator amplification is expected to dampen that of prey (Pompanon et al., 2012; Vestheim and Jarman, 2008). Other approaches are applied to ensure that the samples belong to the relevant species, including the diagnostic PCR which is specific for the expected species and/or Sanger sequencing which reveals only the major DNA sequence of the sample (Bohmann et al., 2011; Forin-Wiart et al., 2018). However, these alternatives add a supplementary step to the process, thereby increasing the time and cost of the analyses. Most importantly, they do not reveal the presence of non-target predator species, or the rate of between-species contamination of samples which – unlike metabarcoding – prevents contaminated samples being discarded. Moreover, we have shown that simultaneous predator amplification by metabarcoding does not necessarily lead to a drop in prey detection. Our results revealed that there is no effect of the percentage of bat reads on the percentage of arthropod taxa detected in mock communities, neither on the number of arthropod occurrences in guano samples. This corroborates the results of Galan et al. (2018) which highlighted the well-balanced proportions of reads for bats and their prey with a particular primer sets. We therefore recommend the systematic identification of predator species when working on environmental faecal samples.

### 4.2 Choosing between the “16S + COI” and “COI alone” strategies

In this study, we have compared primers designed from two mitochondrial genes that are frequently used in eDNA metabarcoding of animals (Deagle et al., 2014): the COI and 16S genes. Our results showed that the 16S primer sets are always less efficient and have lower levels of taxonomic resolution than most of the COI primer sets tested. This was surprising, as for example, the 16S Epp-degen was identified as one of the best primer sets from the *in silico* analysis and we expected a high amplification success due to its short amplicon length and high level of degeneration. In consequence, it is likely that the poor performance of 16S primers revealed from guano analyses is due to the paucity of reference sequences in the 16S database, as has been emphasized in previous studies (Clarke et al., 2014; Elbrecht et al., 2016; Marquina et al., 2019b). This difference in diversity between COI and 16S reference databases could explain the differences in the number of taxa affiliated. Lack of reference sequences could also bias results when analysing short length amplicons. Indeed, as there are less reference sequences, the risk of obtaining an identification with strong confidence levels for the wrong taxa are potentially higher. Moreover, this lack of reference sequences also increases the possibility that 16S affiliations are different from COI ones, therefore leading to false increase in species richness when combining both genes, as it was observed in this study for mock community analyses (Trichoptera-1 identified as *Stenophylax vibex* for COI primer sets and *Anabolia bimaculata* for 16S primer sets; Table S12). Therefore, the choice of the marker in metabarcoding studies should strongly be guided by the comprehensiveness of the reference database. We advocate for the use of the COI gene in animal metabarcoding studies because of its extensive database. However 16S gene can be used if a sequence database of local species of interest is specifically created (Elbrecht et al., 2016).

### 4.3 How many COI primer sets should be used?

To counter the negative effect of primer bias, two main strategies have recently been proposed based on the COI gene. Corse et al. (2019) advocate for the use of multiple primer sets as a key to finely describe species diversity, to reveal greater diversity and to decrease false negative results. However, the *in silico* analysis of the three primer sets used in their study showed that they are not degenerated enough to correctly amplify a large spectrum of prey. Nevertheless, the negative effects of primer bias can be reduced by incorporating primer degeneracy and carefully choosing primer sets suited for the targeted ecosystem and taxonomic groups of interest (Elbrecht et al., 2019). In our study no primer set alone was able to detect all occurrences. This was also the case in the very recent primer set comparison of Elbrecht et al. (2019) on a Malaise trap. At first glance, our results show that combining up to four primer sets allows a gain of between 8.5% (Zeale + fwh1) and 17% (Zeale + fwh1 + MG2 + Epp-degen) of arthropod occurrences in *R. ferrumequinum* samples and between 20.8% (Zeale + fwh1) and 26% (Zeale + fwh1 + MG2 + Epp-degen) of arthropod occurrences in *M. emarginatus* samples, therefore advocating for the use of multiple primers. Deciphering between the ‘one-locus versus several locus strategies hence relies on the trade-off between on one hand the completeness and on the other hand the cost of combining several primers. However, a mean of 23.8% (*R. ferrumequinum*) and 31.6% (*M. emarginatus*) of these occurrences were characterized by the amplification of only two replicates out of three and a small number of reads (median < 125 reads, except Zeale > 4,000 reads in *M. emarginatus* samples). These particular occurrences are the reason why the plateau of taxa occurrence cannot be reached with the combination of a reasonable number of primer sets. These less repeatable occurrences could be due to the weak biomass of prey (low DNA quantity), traces of old meals (degraded DNA, as observed with the Hemiptera-1 of our mock community), traces of secondary predation (e.g. meal of Araneae, the more frequent order in the *M. emarginatus* diet), or environmental contaminations potentially from other bat or insectivorous species. Moreover, 79.3% (*M. emarginatus*) and 86.7% (*R. ferrumequinum*) of the occurrences that were revealed by a single primer set, and therefore that could be interesting to recover by combining several primer sets, are less repeatable between PCR replicates (i.e. only two positive PCRs over three for these taxa; small number of reads). These results bring new insights into the real benefit of attempting to recover all taxa occurrences, in the case where one third of them are less repeatable with regard to PCR replication, and not replicable between primer sets. We therefore recommend the use of a single primer set, following the characteristics described below.

### 4.4 What is the best COI primer set for characterizing insectivorous diets?

*In silico* and *in vivo* tests have been shown to be complementary and critical for assessing the performance of primers (Alberdi et al., 2018; Corse et al., 2019; Elbrecht et al., 2019). Indeed, in our study combining both *in silico*, mock community and guano analyses enabled us to reveal both the strong and weak points of 12 primer sets for a large spectrum of taxa. DNA quality is not of importance during *in silico* and mock community analyses, so that the amplicon length has no significant effect on the number of arthropod taxa detected, while the degeneracy level of the primers has a major positive effect as it minimizes mismatches for DNA from diverse taxonomic assemblages (Braukmann et al., 2019). On the other hand, when considering DNA from guano samples, amplicon length was the most important factor influencing the success of arthropod detection. Too long amplicons (> 313 bp), such as Hex and fwhFol, were less effective, which could be explained by the low proportion of large size DNA fragments in degraded samples. Therefore, our results strongly supported the use of short length amplicon and degenerated primers to maximize biodiversity coverage in metabarcoding analyses (Elbrecht et al., 2019; Elbrecht and Leese, 2017; Galan et al., 2018; Vamos et al., 2017).

Zeale primer set (Zeale et al., 2011) showed important amplification failure in the *in silico* and mock community analyses but not in the guano analysis. Low taxon recovery of the Zeale primer set has previously been observed for terrestrial arthropods (Brandon-Mong et al., 2015; Clarke et al., 2014; Elbrecht et al., 2019). Zeale primer set seems well suited for detecting Lepidoptera because it has a high taxonomic coverage for this order (Clarke et al., 2014; Zeale et al., 2011). As the diet of European bats is mainly dominated by Lepidoptera and Diptera (Alberdi et al., 2019b), the Zeale primer set remains mostly used in insectivorous bat diet studies (e.g. Aizpurua et al., 2018; Andriollo et al., 2019; Clare et al., 2014b; Vesterinen et al., 2018). However, it is not well suited when facing a large spectrum of arthropods. Moreover, it is important to note that the Zeale primer set does not enable to identify bat species. It is thus not recommended in eDNA studies, especially when several insectivorous predator species live on the same sites. Lastly, numerous taxa detected with Zeale primer set are not observed using other primer sets, as noted by Elbrecht et al 2019 who underlined the putative presence of false positive results, or are weakly reliable considering PCR replication results. Therefore, previous results of insectivorous (bat) diet gathered from this primer set only might be affected by the potential biases revealed here.

Here, the best primer set identified to characterize simultaneously the diet of *R. ferrumequinum* and *M. emarginatus* from guano samples collected under mixed roosts was fwh1 (Vamos et al., 2017) (Table 2). The contrasted performances of the primer sets between bat species for arthropod detection highlight the need to test several primer sets on a representative subset of eDNA samples before undergoing large-scale metabarcoding study in order to determine which is the most suitable for the sampling scheme and biological model.

## 5 Conclusion

Our study supports the importance of combining *in silico*, mock communities and field sample preliminary analyses to determine the benefits and the limits of the potential primer sets before conducting research based on metabarcoding. This three-stage assessment of primer performance confirms that primer success is determined by amplicon length, base degeneracy level and completeness of reference databases. Our work also emphasizes that the identification of the best primer sets for insectivorous diet studies is not only highly dependent on the objective and financial resources of the study, but also on the sampling scheme and fieldwork constraints that impact DNA quality and the need to identify the predator species. Finally, we have evidenced the presence of unreliable occurrences that significantly reduces the benefits of using combinations of primer sets. In conclusion, we advocate for the use of multi-criteria assessment that summarizes all the information required to evaluate primer sets performance to guide the choice of the investigator. This analytical framework can easily be adapted to other metabarcoding studies of predator diet.

## ACKNOWLEDGMENTS

We are very grateful to A. Cheron, J. Dechartres, M. Dorfiac, Y. Prioul, J.B. Pons and all the people involved in the guano and patagium sampling. We also thank L. Benoit, A. Foucart, L. Soldati, M. Sorel and J-C. Streito for the arthropod sampling and identification, V. Elbrecht for the COI sequence alignments, J-F. Martin for the dual-indices, F. Dorkeld for the *blastn* script used for the 16S taxonomic assignment. Financial support was received from the LABEX ECOFECT (ANR-11-LABX-0048) of Université de Lyon, within the program “Investissements d’Avenir” (ANR-11-IDEX-0007) operated by the French National Research Agency (ANR), from the Institut National de la Recherche Agronomique (EFPA department) and the internal funding from the CBGP laboratory. O. Tournayre PhD is funded by the LabEx CeMEB, an ANR “Investissements d’Avenir” program (ANR-10-LABX-04-01). Data used in this work were partly produced through the genotyping and sequencing facilities of ISEM (Institut des Sciences de l’Evolution-Montpellier) and LabEx Centre Méditerraneen Environnement Biodiversité.

## DATA ACCESSIBILITY

Supplementary data deposited in Dryad (https://doi.org/10.5061/dryad.4v9n227) include: (i) raw sequence reads (fastq format), (ii) raw abundance tables and (iii) reference sequences of the 33 arthropod taxa and the bat used in the mock communities.

## AUTHORS CONTRIBUTION

The study was conceived and designed by M.G., O.T., D.P. and N.C. The sampling schemes were designed by M.L., M.G., O.T., O.F.C., D.P., N.C., and conducted by M.L., O.F.C., O.T. and M.G. Laboratory protocols were designed and performed by M.G. and O.T. S.P., M.G., O.T. and M.T. performed the bioinformatics analyses and O.T. carried out the statistical analyses. A first draft of the manuscript was written by O.T, M.G., D.P. and N.C. All authors contributed to the writing of the final version of this paper.

## SUPPORTING INFORMATION

**Appendix S1.** Rules used to determine the final identifications of OTUs in guano samples.

**Table S1.** Number of COI and 16S OTUs, obtained respectively with sequences from BOLD and NCBI databases.

**Table S2.** Information about the samples, the laboratory controls and the technical replicates.

**Table S3.** Versions of the primers used in PCR_1_ of the 2-step PCR including heterogeneity spacers and Illumina sequencing primer sequences.

**Table S4.** Information about the samples, the laboratory controls and the technical replicates. Error-proof indexes for high throughput sequencing were created by Martin (2019).

**Table S5.** Genuine sequences of the two mock communities. Number of reads are reported for each replicate and each primer set.

**Table S6.** Examples of affiliation errors for several primer sets in reference databases.

**Table S7.** Detailed criteria and results for the *in silico* and empirical comparison of the twelve primer sets.

**Table S8.** Mean penalty scores obtained for each arthropod order (OTUs > 100) with each primer set using *PrimerMiner* program.

**Table S9.** *In silico* amplification success obtained for each arthropod order (OTUs > 100) and primer set using *PrimerMiner* program.

**Table S10.** Effect of the primer set characteristics on the in silico amplification success and mean penalty score, the detection of mock community taxa and the detection of arthropod occurrences in guano samples. *** p < 0.001, ** p < 0.01 and * p < 0.05.

**Table S11.** Objectives and impact of the pre-process and FROGS pipelines on the number of reads. ¶ ‘Complete run ‘ indicates that the run included other samples than those of the two mock communities. ¥ = genuine sequences, † = identity > 97% and coverage > 90%, ‡ identity < 97% and/or coverage <90%.

**Table S12.** Molecular identification of the taxa in the mock communities obtained using the genuine sequences of each specimen. ‘<97%’ indicated a percentage of identity lower than 97%. Table S13. Percentage of each bat species in mixed samples. Rf = *R. ferrumequinum,* Me *= M. emarginatus.*

**Figure S1.** Consensus sequence alignment of 21 arthropod orders using the *PrimerMiner* R package.

**Figure S2.** PCR products of the mock communities (left column) and guano samples (right column) on agarose gel.

**Figure S3.** Percentage of bat reads for each primer set in the mock community MC_arthr+bat_. The red line indicated the expected value for percentage of bat reads (50%).

**Figure S4.** Representation on a log-scale of the number of reads gathered for each arthropod order with the 12 primer sets, from the three technical replicates of the mock community MC_arthr_ and MC_arthr+bat_. The blue line indicates the expected number of reads. Each point represents a technical replicate. Yellow bars emphasize situations where only one or two PCR over three amplified. Red bars emphasize situations where none of the replicate amplified.

**Figure S5.** Percentage of bat reads gathered in the guano samples (N_Rf_ = 10, N_Me_=11) for each primer set. *Me = Myotis emarginatus,* Rf *= Rhinolophus ferrumequinum.*

**Figure S6.** Occurrences of taxa in *M. emarginatus* samples (N = 11) and *R. ferrumequinum* (N = 10) not shared between primer sets, or shared by at least one COI primer set and one 16S primer set, or shared by at least two COI primer sets. Number of PCR replicates: ‘2’ (light green) = occurrence in two PCR over three, ‘3’ (green) = occurrence in three PCR over three.

