## Supplementary figures and images for "*In silico* and empirical evaluation of twelve COI & 16S metabarcoding primer sets for insectivorous diet analyses"

### Figure S1

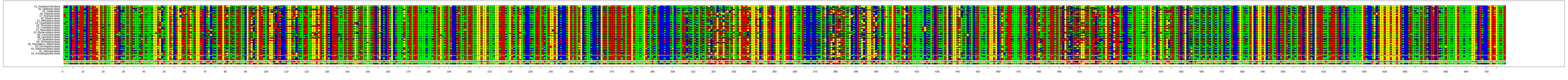
