## Appendix S1 for "*In silico* and empirical evaluation of twelve COI & 16S metabarcoding primer sets for insectivorous diet analyses"

**Appendix S1.** Rules used to determine the final identifications of OTUs in guano samples.

1. The identity percentage must be ≥ 97% following the confidence levels described in Galan et al. (2018) and the coverage percentage must be ≥ 90 %. The BLAST e-value must be < 1.10^-40^.
2. For each sample, if several OTUs have the same assignment and confidence level, we only kept the one with the highest number of reads.
3. We removed OTUs corresponding to orders that bats do not intentionally eat (bacteria, fungi, rotifera, nematoda, branchiopoda, siphonaptera, mesostigmata). These orders most likely corresponded to environmental contaminations or OTUs from bat or arthropod gut. We also discarded OTUs showing putative assignment problems, i.e. that were assigned to species or genus that are not expected in France and in Europe (according to the Fauna Europaea database (Jong et al., 2014) and the Inventaire National du patrimoine Naturel (INPN ; Muséum national d’Histoire naturelle, 2003)).
4. In case of pseudogene/chimera suspicion - very close species in the same sample with high differences in number of reads and slight differences in confidence criteria -, we only kept the species with the highest number of reads and the highest confidence criteria.
5. We manually checked every remaining OTUs in BOLD and GENBANK to identify errors (see Table S6 for several examples). Suspicious sequences were checked in the BioEdit software (Hall, 1999) to detect potential remaining chimera.
6. Finally, we clustered the arthropods that exhibited slight assignment differences within a given sample when considering several primer sets. This clustering was done following three conditions: the arthropods shared a common family assignment rank, had a high number of reads in the sample, and we observed the same pattern in several samples. We decided to apply this last step because of (i) differences in resolution levels between primer sets (short vs. long fragments) and gene (16S - COI), (ii) presence of species complex for which both genes were not resolutive (e.g. *Tipula sp.*), and (iii) potential biases in reference databases between genes (16S *vs* COI) and between primer sets (at the beginning *vs* at the end of the COI).
